# Common and distinct genetic architecture of age at diagnosis of diabetes in South Indian and European populations

**DOI:** 10.1101/2022.09.14.508063

**Authors:** Sundararajan Srinivasan, Samuel Liju, Natarajan Sathish, Moneeza K Siddiqui, Ranjit Mohan Anjana, Ewan R. Pearson, Alexander S.F. Doney, Viswanathan Mohan, Venkatesan Radha, Colin N.A Palmer

**Author notes:** **ADDRESS FOR CORRESPONDENCE:** Professor Colin NA. Palmer,. Division of Population Health and Genomics, Ninewells Hospital and Medical School, University of Dundee, United Kingdom.

## Abstract

South Asians are diagnosed with type 2 diabetes (T2D) more than a decade earlier in life than seen in European populations. We hypothesised that studying the genomics of age of onset in these populations may give insight into earlier age at onset of T2D among individuals of South Asian descent. We conducted a meta-analysis of GWAS of age at diagnosis of T2D in 34,001 individuals from four independent cohorts of European and Asian Indians. We identified two signals near the *TCF7L2* and *CDKAL1* associated with age at onset of T2D. The strongest genome-wide significant variants at chromosome 10q25.3 in *TCF7L2* (lead SNP rs7903146; p = 2.4 ×10^-12^; p-het =0.01; Beta = -0.436; SE = 0.02) and chromosome 6 p22.3 in *CDKAL1* (rs9368219; p = 2.29 ×10^-8^; p-het =0.007; Beta = -0.053; SE=0.01) were directionally consistent across ethnic groups and present at similar frequencies, however both loci harboured additional independent signals that were only present in the South Asian cohorts. A genome wide signal was also obtained at chromosome 10q26.12 in *WDR11* (rs3011366; p = 2.4 ×10^-8^; p-het =0.25; Beta = 1.44; SE=0.25) specifically in the South Asian Indian cohorts. Our study estimated 17% of heritability (h^2^) for age onset of T2D among South Indians; higher than the 5% heritability we observed in Europeans suggesting different genetic architectures between these populations. We further estimated the Asian Indian genome-wide polygenic risk score for T2D risk and identified that the variance explained by the Indian specific polygenic risk score (PRS) for age at onset of T2D is substantially higher (2%) in an independent cohort of Asian Indians compared to that seen in White Europeans (<0.1%). This reveals that although variants do exist that are shared between the two population there are also genetic disparities of age onset of T2D between the ethnicities.

**Author Summary:** Recent large multi-ancestry genome-wide meta-analyses have identified over 277 genetic loci associated with type 2 diabetes (T2D). Most association studies for T2D have focused on populations of European ancestry, and the specific genetic architecture of T2D in the South Asian population has not been extensively investigated. Increasing evidence indicates that the prevalence of T2D is higher in South Asians with a strikingly earlier age at diagnosis when compared to Europeans. Genome-wide association studies (GWAS) on age at age at diagnosis of T2D are limited and can help elucidate underlying mechanisms. We conducted the largest meta-analysis to date of genome-wide association studies of age at diagnosis as a surrogate for age of onset of T2D in South Indians and Europeans. Our trans-ancestry meta-analysis identified genetic loci near *TCF7L2* and *CDKAL1* with substantial ethnic heterogeneity in both allele frequencies and effect sizes in early diagnosis with T2D. Two novel variants near *TCF7L2* (rs570193324) and *CDKAL1* (rs143316471) were associated with age of diagnosis of T2D only in South Indians and were present at much lower frequency in the European populations. Heritability estimates for age of onset were much stronger in Asian Indians compared to Europeans and a polygenic risk score was constructed using a South Indians, which explained about 2% trait variance compared to European ancestry (<0.1%). Our novel findings provide a better understanding of ethnic differences in the age at onset of T2D and indicate the potential importance of ethnic differences in the genetic architecture underpinning T2D.

## Introduction

Type 2 diabetes (T2D) is a multifactorial disease characterised by impaired insulin action and pancreatic islet dysfunction. The global prevalence of T2D is a pivotal driver of cardiovascular and renal disease[1–3] affecting hundreds of millions of people globally and is responsible for long-term complications, decreased quality of life, and increased mortality[4–7]. Improved understanding the intrinsic genomic and phenotypic heterogeneity driving T2D has major potential for improvement of its clinical management and reducing morbidity and mortality. South Asian Indians have an earlier age of onset of diabetes compared to Europeans and mounting evidence suggests this is associated with earlier mortality emphasising the need to delay or prevent the onset of T2D in this ethnic group[8,9]. South Asians with newly diagnosed diabetes may have a higher risk for microvascular complications than Europeans[10]. Recent studies highlight that higher cardiovascular mortality and disease risk is associated with early onset of T2D diagnosis compared to individuals with delayed onset of the disease [11]. South Asians (individuals originating from India, Pakistan, and Bangladesh) are genetically more diverse than the white Europeans, and the prevalence of T2D is much higher in this ethnic group than other ethnic backgrounds [14–16].

Currently nearly 250 genetic loci (more than 400 unique genetic variants) have been identified that influence T2D[2,12,13]. Several of these genetic loci have only been identified in European study populations. A trans-ethnic meta-analysis of European and East Asian populations reported several T2D risk variants with significant allelic frequency heterogeneity[14]. Such frequency differences between ethnic populations affects the power to detect genomic signals within a specific ethnic subgroup. A recent study reported that migrant South Asians are more insulin resistant and have poorer β-cell function at a younger age than White Europeans [17]. Previously identified genetic variants explained about (~10%) of the heritability of T2D [15].

Despite advancement in genetic research tools, South Indian specific studies are minimal compared to European ancestry. To our knowledge, no GWAS has been published that addresses the age at diagnosis of T2D in people of South Asian Indian ethnicity and compare this with European populations. We aimed to identify novel genetic determinants that influence the risk of younger age of diagnosis in two distinct ethnic backgrounds more specifically in South Asians and Europeans. We aimed to develop, evaluate, and understand a T2D age at diagnosis PRS in Asian Indians (DMDSC) and Europeans (GoSHARE) used in the study cohort. In this multicentre study, we focused on inter-ancestry differences in the genetics of age at onset of T2D that might influence ethnic-ancestry differences in health outcomes in general and T2D.

## Results

A total of 34,001 participants with T2D were included for this study after-quality control filtering: 8,295 T2D patients of Asian Indians ancestry from a large tertiary diabetes centre, Dr Mohan’s Diabetes Specialties Centre (DMDSC), 6,999 patients of European ancestry from Genetics of Diabetes Audit and Research Tayside and Scotland (GoDARTS)[16], 4,155 patients of European ancestry from Scottish Health Research Register (SHARE)[17] and 14,552 T2D participants from United Kingdom Biobank (UKBB). We identified participants of European (N=13,744) and South Asian Indians (808) in the UKBB using principal components (PCs) analysis of genome-wide data and found that this was consistent with self-reported ancestry information (Supplement Fig 1). A detailed illustration of the study design is presented in Fig 1. The population characteristics of the cohorts are described in Table 1. Notably, we observed the average age of diagnosis of T2D in the Asian Indians was 40 years, whereas in the white Europeans, the mean age of diagnosis was 60 years.

**Fig 1.**
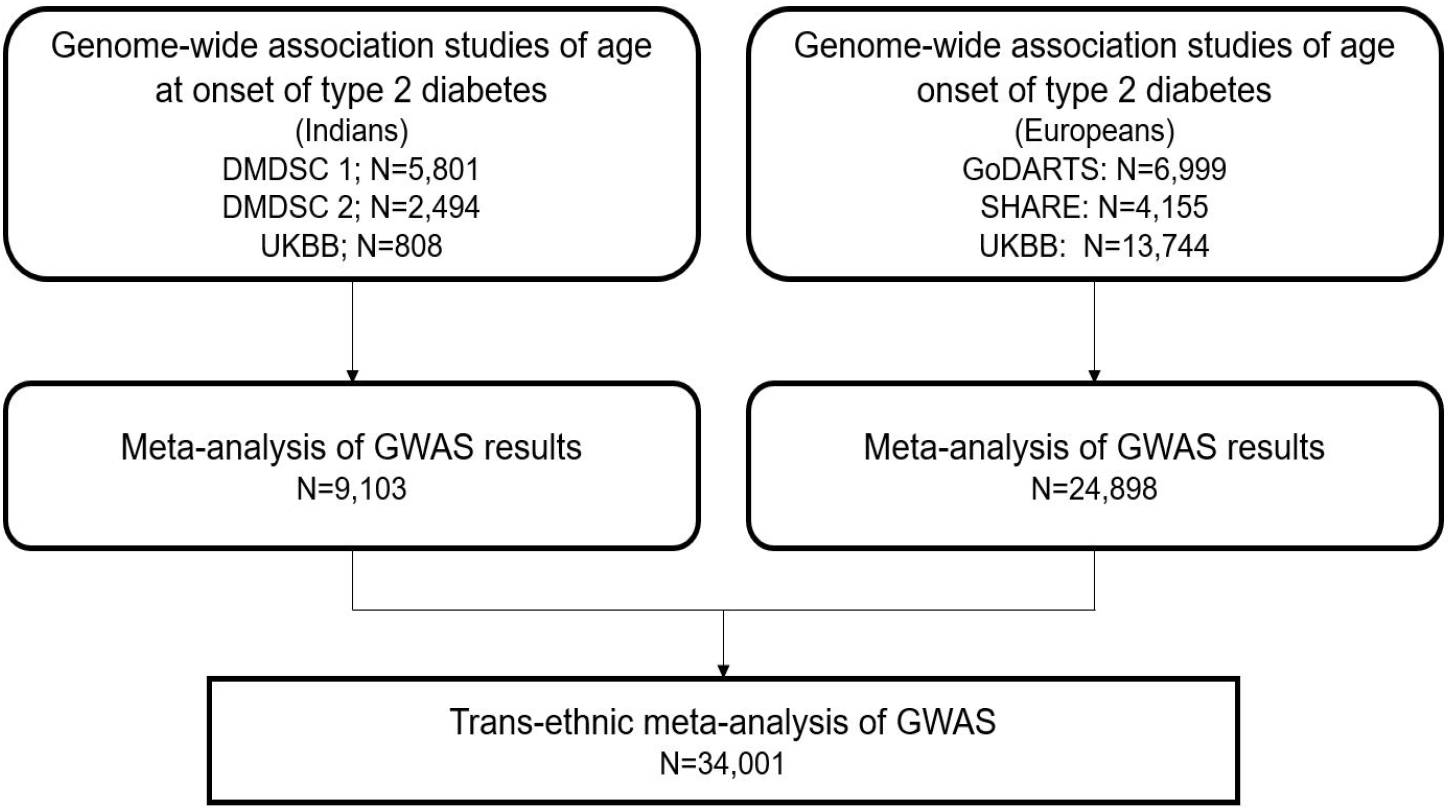
Study design.

**Table 1.**
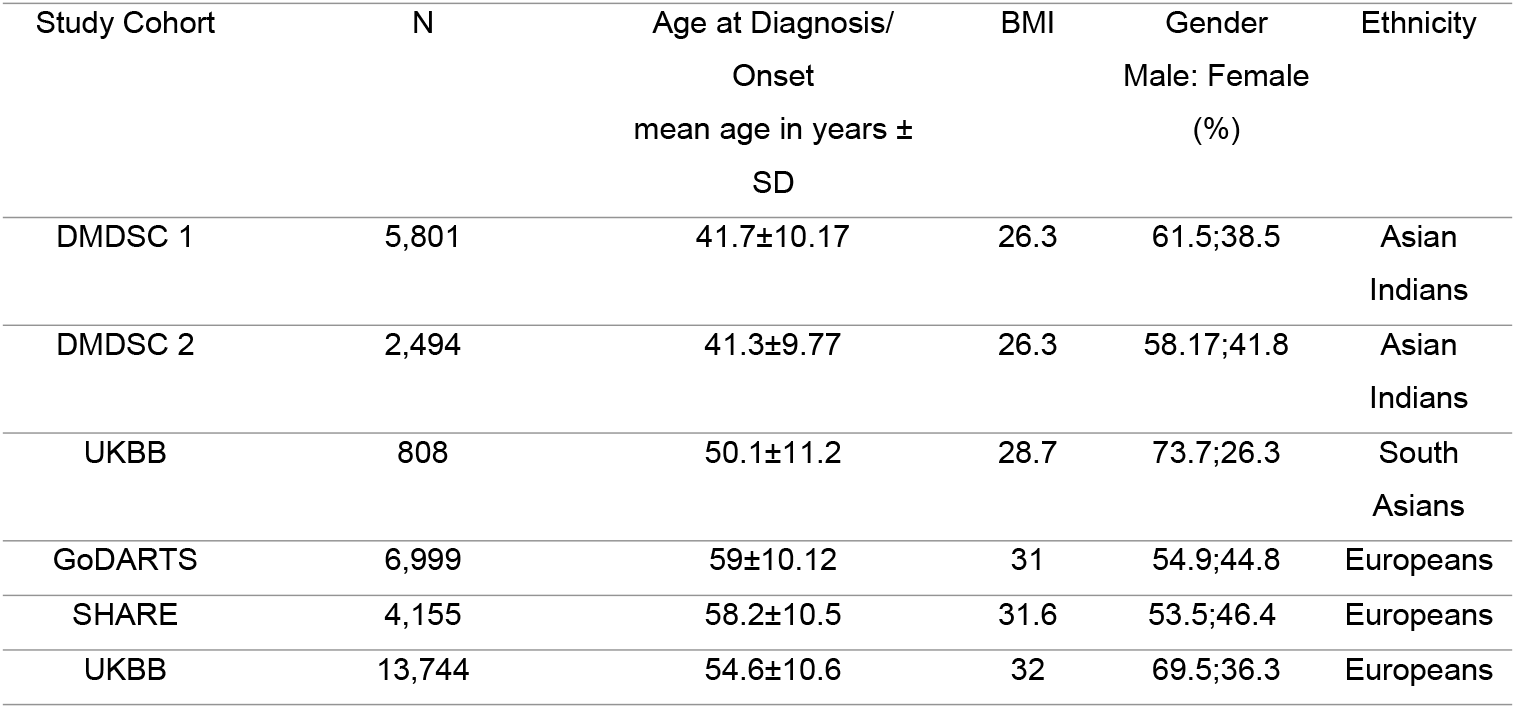
Characteristics of the study population.

| Study Cohort | N | Age at Diagnosis/<br>Onset<br>mean age in years ±<br>SD | BMI | Gender<br>Male: Female<br>(%) | Ethnicity |
| --- | --- | --- | --- | --- | --- |
| DMDSC 1 | 5,801 | 41.7±10.17 | 26.3 | 61.5;38.5 | Asian<br>Indians |
| DMDSC 2 | 2,494 | 41.3±9.77 | 26.3 | 58.17;41.8 | Asian<br>Indians |
| UKBB | 808 | 50.1±11.2 | 28.7 | 73.7;26.3 | South<br>Asians |
| GoDARTS | 6,999 | 59±10.12 | 31 | 54.9;44.8 | Europeans |
| SHARE | 4,155 | 58.2±10.5 | 31.6 | 53.5;46.4 | Europeans |
| UKBB | 13,744 | 54.6±10.6 | 32 | 69.5;36.3 | Europeans |

### SNP-based heritability

Using the LDSC tools, we estimated the SNP-based heritability for Age onset of T2D in Asian Indians was 17% (SE 6%) but was only 5% (SE 2%) for Europeans.

### Trans-ethnic Meta-analysis of GWAS for age of T2D diagnosis

The genome-wide association analyses of age at diagnosis of type 2 diabetes were conducted for each cohort separately using a linear mixed model as implemented in BOLT-LMM. We then conducted trans-ancestry meta-analyses by utilising Haplotype Reference Consortium (HRC) imputed data up to 26.2 million SNPs directly genotyped or successfully imputed at high quality across all the data sets. Association statistics and P values for heterogeneity were combined using fixed-effects meta-analyses as implemented in METAL [18]. Our meta-analysis revealed two previously known T2D loci at chromosome 10 q25.2 near Transcription factor 7-like 2 (rs79603146, *TCF7L2*, P < 2.4 x 10^-12^, Beta -0.436; SE = 0.02; P-het =0.01) and at chromosome 6p22.3 cyclin-dependent kinase 5 (*CDK5*) regulatory subunit-associated protein 1-like 1 (*CDKAL1*) (rs9368219, P < 2.29 x 10^-8^, Beta -0.053; SE 0.01; P-het =0.007) associated with T2D age-diagnosis (Fig 2). The allelic frequency of CDKAL1 is more common in DMDSC Asian Indian cohort (MAF = 0.26) compared to Caucasians (MAF =0.18) The lead SNPs at *TCF7L2* and *CDKAL1* (Fig 3A and 4A) locus demonstrated consistent allelic direction across all cohorts, with the risk alleles associated with lower age of diagnosis, however, a large difference was observed in the size of the estimate of the effects between the South Indian and European cohorts explaining that variation in allelic effect estimates is presumably due to their genetic ancestry it being. Interestingly the effect size of the variants was much lower in the cohorts of European descent. Ethnic-specific meta-analysis results are presented in supplement tables.

**Fig 2.**
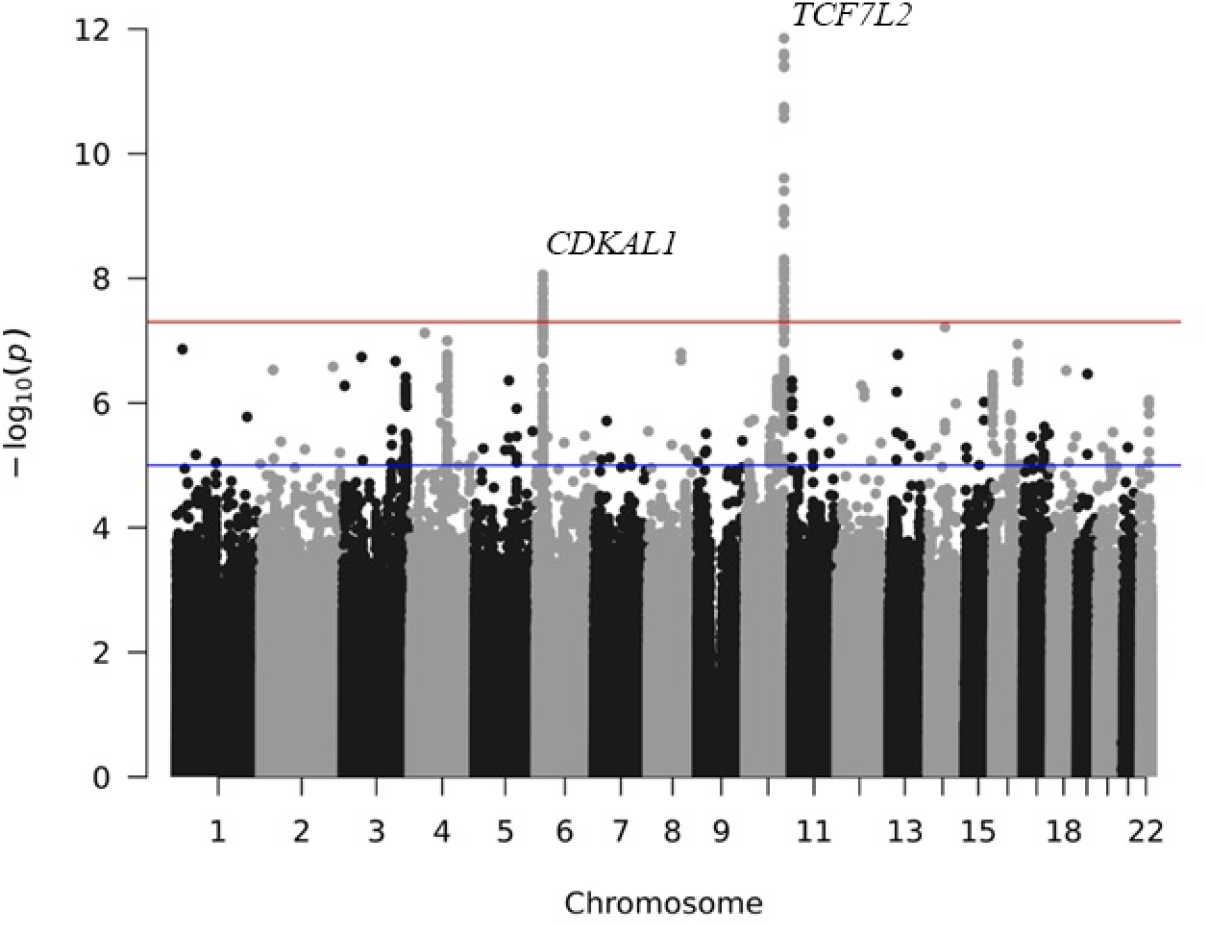
Manhattan plot showing the P-value of association tests for SNPs with Age at diagnosis of T2D in a Trans ethnic meta-analysis of GWAS. Two horizontal lines from the bottom indicate the suggestive (P<5×10^-5^) and genome-wide significance threshold (P<5×10^-8^), respectively. The X-axis represents the physical position and the 22 autosomal chromosomes; Y-axis represents the negative logarithm of association p-value. Each dot on the plot represents millions of imputed SNPs across the whole genome.

**Fig 3.**
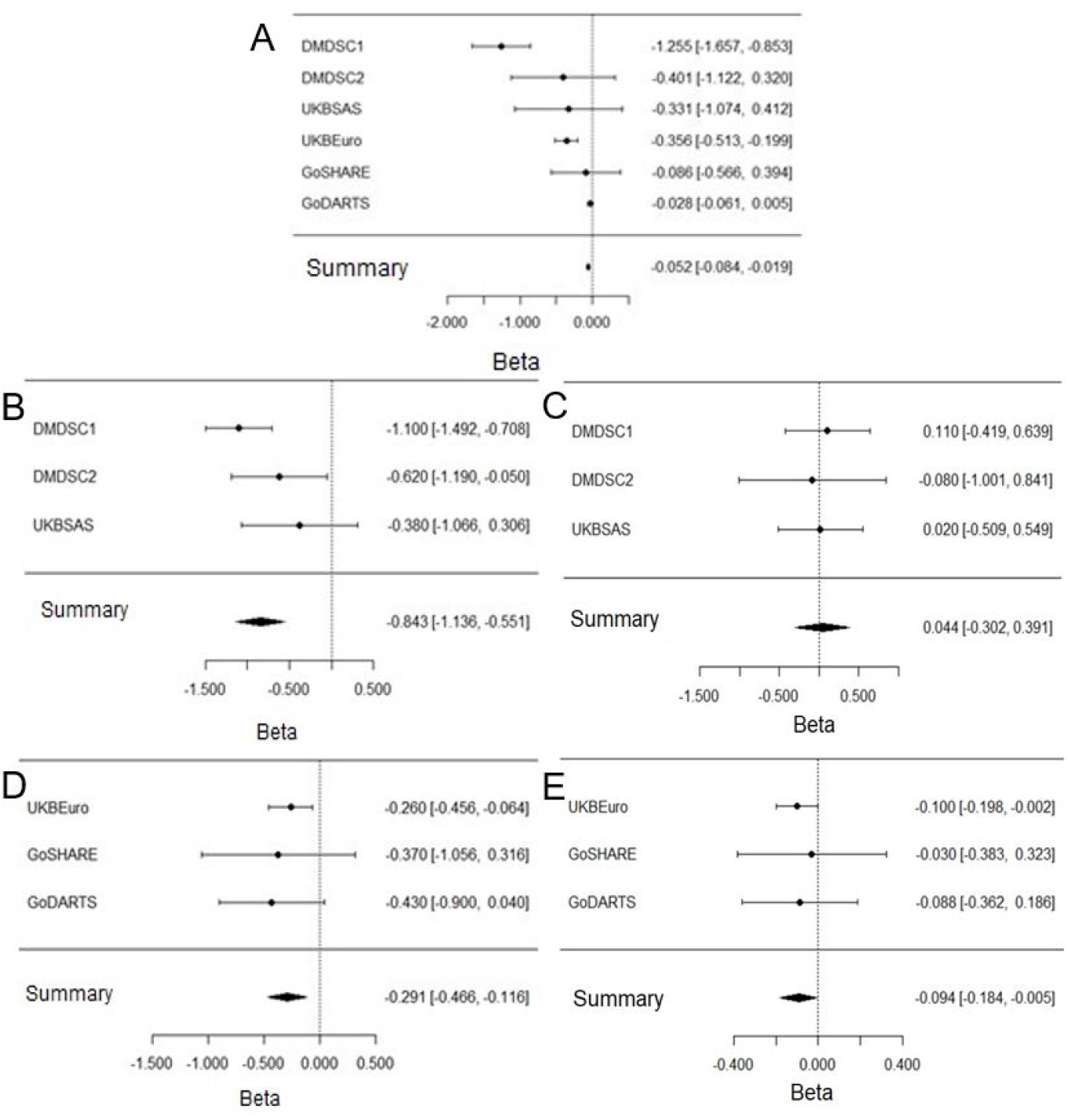
Forest plot for the top significant SNP rs7903146 near *TCF7L2* (Effect allele - T) in individuals with T2D. DMDSC Dr. Mohan Diabetes speciality Clinic, UKBSAS UK Biobank South Asians. GoDARTS, Genetics of Diabetes Audit and Research in Tayside Scotland; GoSHARE, Genetics of Scottish Health Research Register; UKBB Euro, United Kingdom Biobank Europeans. A) Overall meta-analysis of GWAS of Age onset of T2D; B) South Asians with earlier onset of T2D (Age at diagnosis between 20 – 55 years); C) South Asians with later onset of T2D (Age at diagnosis between 20 – 55 years); D) Caucasians with earlier onset of T2D (Age at diagnosis between 20 – 55 years); E) Caucasians with later onset of T2D

**Fig 4.**
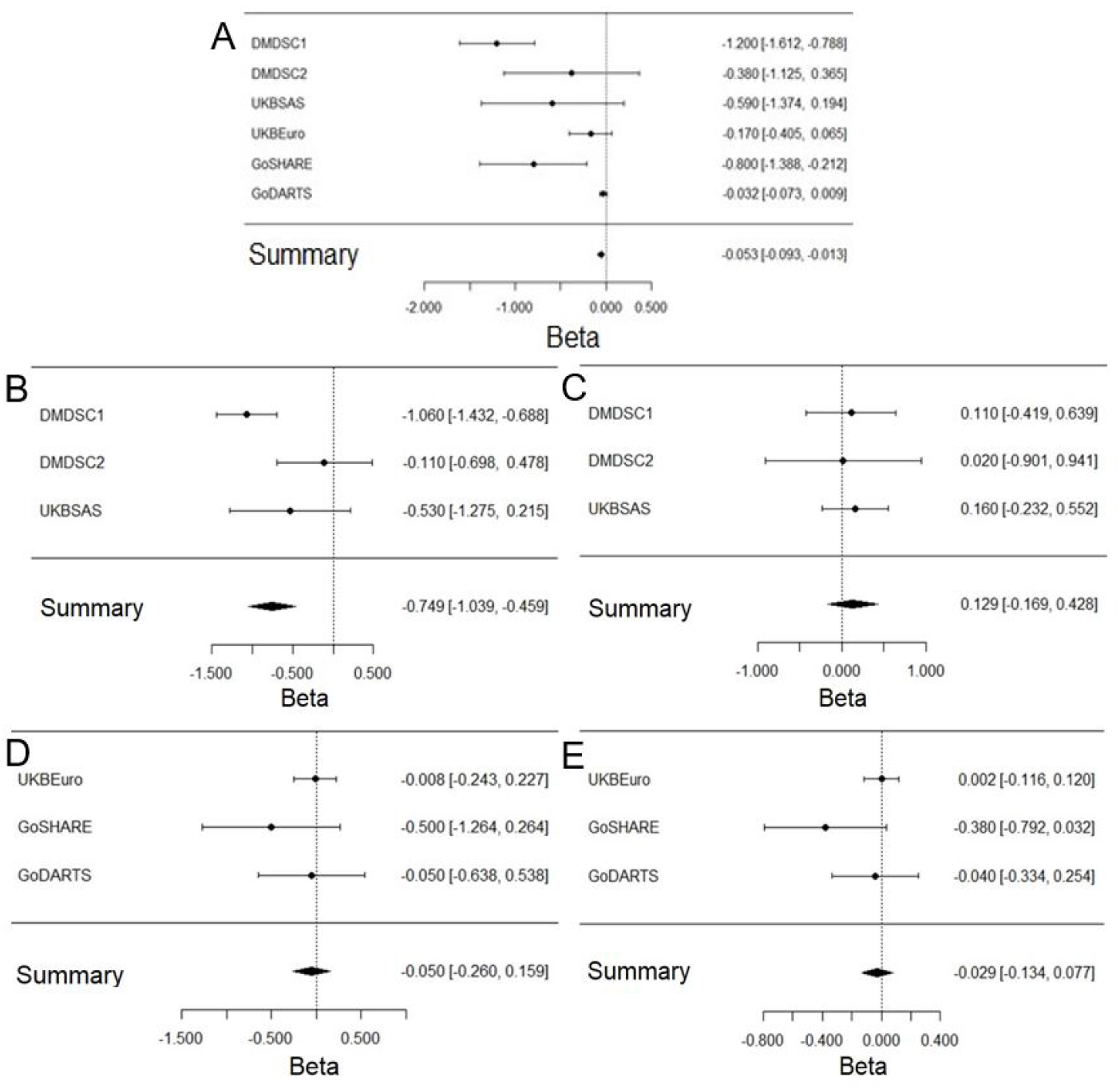
Forest plot for the top significant SNP rs9368219 near *CDKAL1* (Effect allele - T) in individuals with T2D. DMDSC Dr. Mohan Diabetes speciality Clinic, UKBSAS UK Biobank South Asians. GoDARTS, Genetics of Diabetes Audit and Research in Tayside Scotland; GoSHARE, Genetics of Scottish Health Research Register; UKBB Euro, United Kingdom Biobank Europeans. A) Overall meta-analysis of GWAS of Age onset of T2D; B) South Asians with earlier onset of T2D (Age at diagnosis between 20 – 55 years); C) South Asians with later onset of T2D (Age at diagnosis between 20 – 55 years); D) Caucasians with earlier onset of T2D (Age at diagnosis between 20 – 55 years); E) Caucasians with later onset of T2D

### Stratification of Type 2 Diabetes by Age Onset

As the two ethnic groups were very different in the mean age of diagnosis, we explored the extent to which the observed differences in allelic effect size may be determined by the heterogeneity in age of onset between the populations, therefore stratified by of age of onset.

Based on the South Indian mean age of diagnosis (Table 1), the study participants in both ethnicities were stratified into the early-diagnosis T2D group (20-55 years) (Fig 3B & D) and late diagnosis T2D (diagnosed after 55 years) (Fig 3C & E). We found that the effect size of both the *TCF7L2* and *CDKAL1* variants was more pronounced in the early onset group regardless of ethnicity. (Fig 3B, D & 4B, D). These variants have very little effect on age of diagnosis in those with diabetes diagnosed after 55 years of age in either ethnicity (Fig 3C, E& 4C, E).

### The role of other T2D variants in Age of diagnosis

We identified several previously reported T2D variants as suggestive signals (P < 1 × 10^-5^) in this age onset of T2D trans-ethnic meta-analyses (Supplement Table). In particular, the risk variant nearby *SEC24B* at chromosome location 4q25 (rs76170449, P< 1.79×10^-7^) is also associated with cardiovascular traits, and 3p24.3 *ZNF385D* (rs17011243, P< 1.13×10^-5^) associated with T2D in prior GWAS studies. In addition to the other suggestive signals, we detected potential common variants at chromosome location 16p13.3 (rs1977100, *TPSD1*, P < 3.40 × 10^-6^) and 17q21.2 (rs684214, *MLX*, P< 2.40 ×10^-6^) with no difference in their effect estimates between two distinct ethnic groups (Supplement Table). We replicated previously reported South Asian T2D genome-wide signals[19–22] with suggestive evidence or a nominal association for age onset of T2D in the trans ethnic meta-analyses and ancestry-specific groups. Most of the formerly associated T2D loci from earlier GWAS showed consistent effect estimates in South Indian and European subjects. These include *LPL, SLC30A8, GCKR, THADA, HNF1A, TPCN2, GRB14, SIX3, WDR11, SPC25, CENTD2, MLX, APS32, WFS1, ST6GAL1, KNCQ1 and IGF2BP2*.

### Meta-analysis of Asian Indian cohorts

In the meta-analysis of only the Asian Indian cohorts, we also found an additional novel genome-wide signal at chromosome 10 q26.12 near the *WDR11* region (rs3011366, P < 2.46 ×10^-8^, Beta 1.44, SE 0.25). However, this variant was not associated the age-of-onset in the European cohorts (Table 2). WDR11 encodes the WD repeat domain family, which involves signal transduction and cell cycle progression. Previous GWAS studies in the European populations and UK Biobank T2D participants have reported that *WDR11* (rs3011366) was associated primarily with fasting glucose[23]. The Conditional analyses conducted in South Indian ancestry (Table 3), indicated two independent secondary signals at *TCF7L2* (rs570193324, q25.2, P < 3.2E-05, Beta 9.8, MAF 0.002, R2 0.0006) and *CDKAL1* (rs143316471, P < 0.0054, Beta -5.3, MAF 0.003, R2 0.005). Allelic frequency for both independent signals were rare in European cohorts compared to South Indian Cohorts. The regional plot for independent association of *TCF7L2* and *CDKAL1* is shown in Fig 5.

**Table 2.**
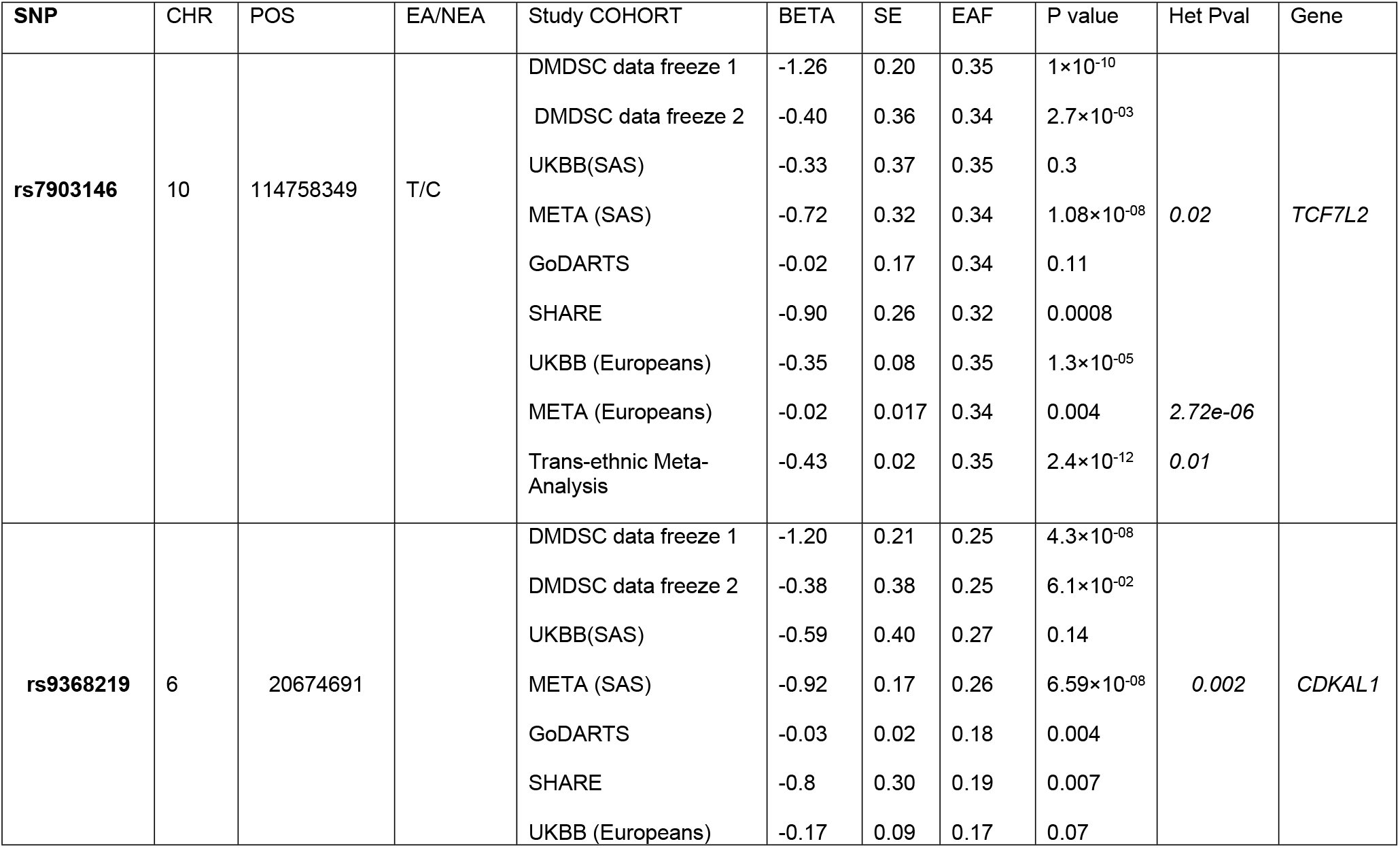
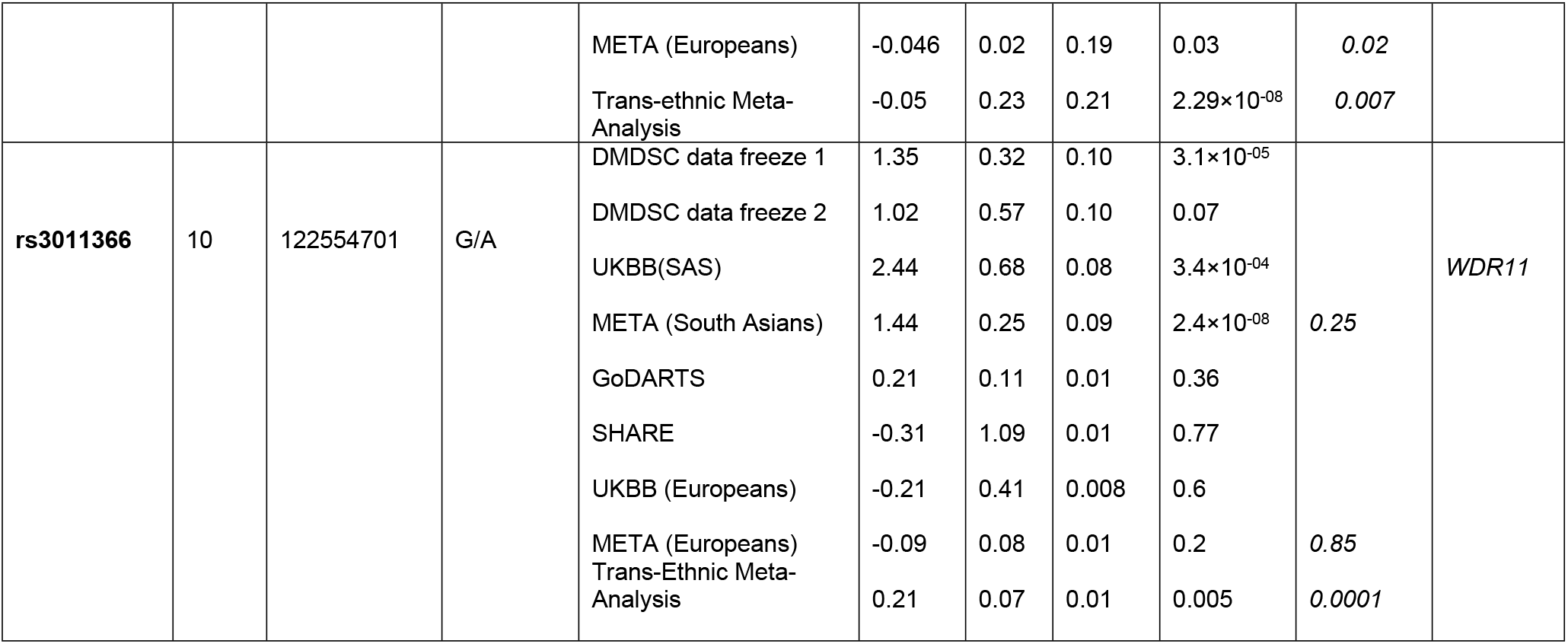
Summary statistics of the most significant SNPs from meta-analysis.

**Table 3.** Summary statistics of the novel variants after conditioning on index SNPs.

| **SNP** | **CHR** | **POS** | **EA/NEA** | **BETA** | **SE** | **P value** | **EAF** | **Gene** |
| --- | --- | --- | --- | --- | --- | --- | --- | --- |
| **rs570193324** | 10 | 114925065 | T/C | 9.8 | 2.3 | 3.2e-05 | *0.002* | *TCF7L2* |
| **rs143316471** | 06 | 20693278 | T/C | -5.3 | 2.3 | 0.0054 | *0.003* | *CDKAL1* |

**Fig 5.**
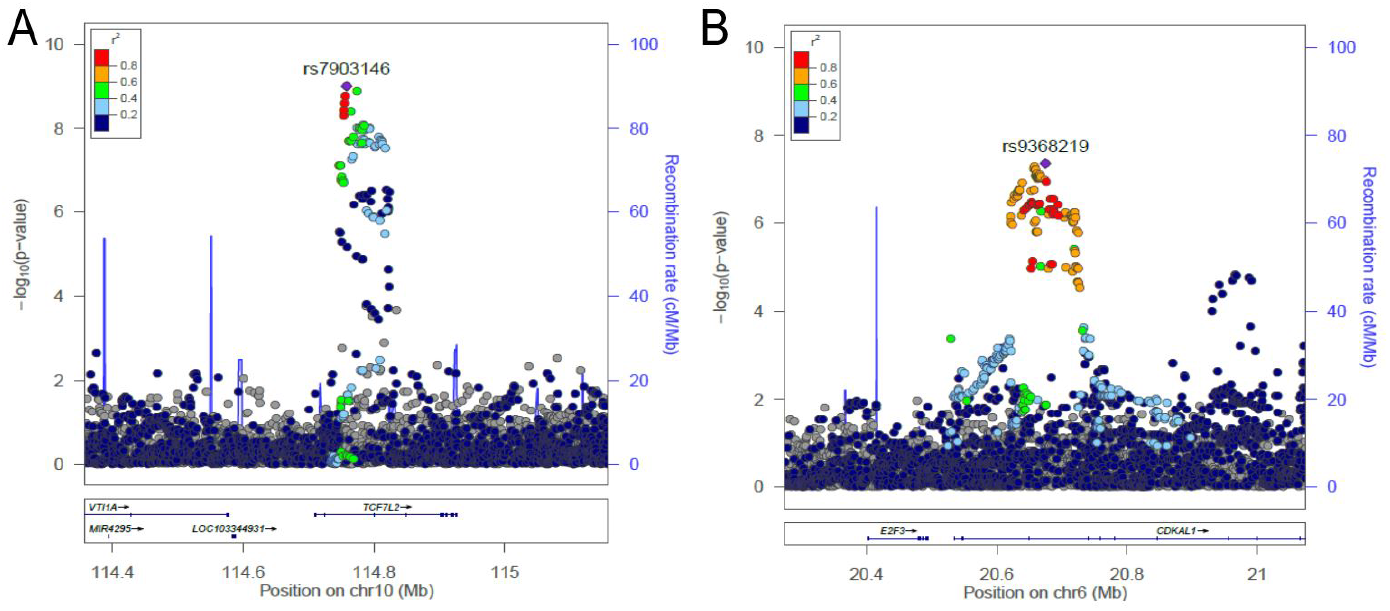
Regional association plots of top significant SNPs. Associations at *TCF7L2* (A) and *CDKAL1* (B) in the South Asian meta–genome-wide association studies. Chr, chromosome; cM/Mb, centimorgan/megabase (genomic location in reference build 37 [Hg19]).

### Meta-analysis of European cohorts

In the analyses unique to White Europeans, we did not observe any genome-wide signal in the European specific meta-analyses. However, we observed suggestive association of a missense variant rs2232328 near *SPC25*, an established variant for fasting blood glucose and type 2 diabetes[24]. Several other SNPs reached suggestive significance for age onset of T2D, and the direction of effect was consistent across all cohorts of European descent (*Supplementary Table*).

### PRS analysis reveals polygenic effects for Age at the onset of Type 2 Diabetes

Polygenic Risk Scores (PRS) are emerging as more informative clinical screening and prediction tool with an increasing number of robust genomic variants identified through more extensive genetic association studies[25]. To investigate whether different genetic variants shared between ethnicities were conferring the risk of the onset of T2D: first, we constructed a PRS for all genome wide significant SNPs associated with age of diagnosis using the DMDSC 1 cohort of the Asian Indian genetic data (n=5801) and validated this using the DMDSC 2 of Asian Indian data which contained no overlapping participants; next we assessed the performance of this South Asian GRS in the European cohort (GoSHARE) (Fig 6). The PRS replicated strongly between the Asian Indian cohorts. On the other hand, the Asian Indian derived PRS explained less than 0.1%of the variance in age of diagnosis of T2D in the GoSHARE cohort (European ancestry).

**Fig 6.**
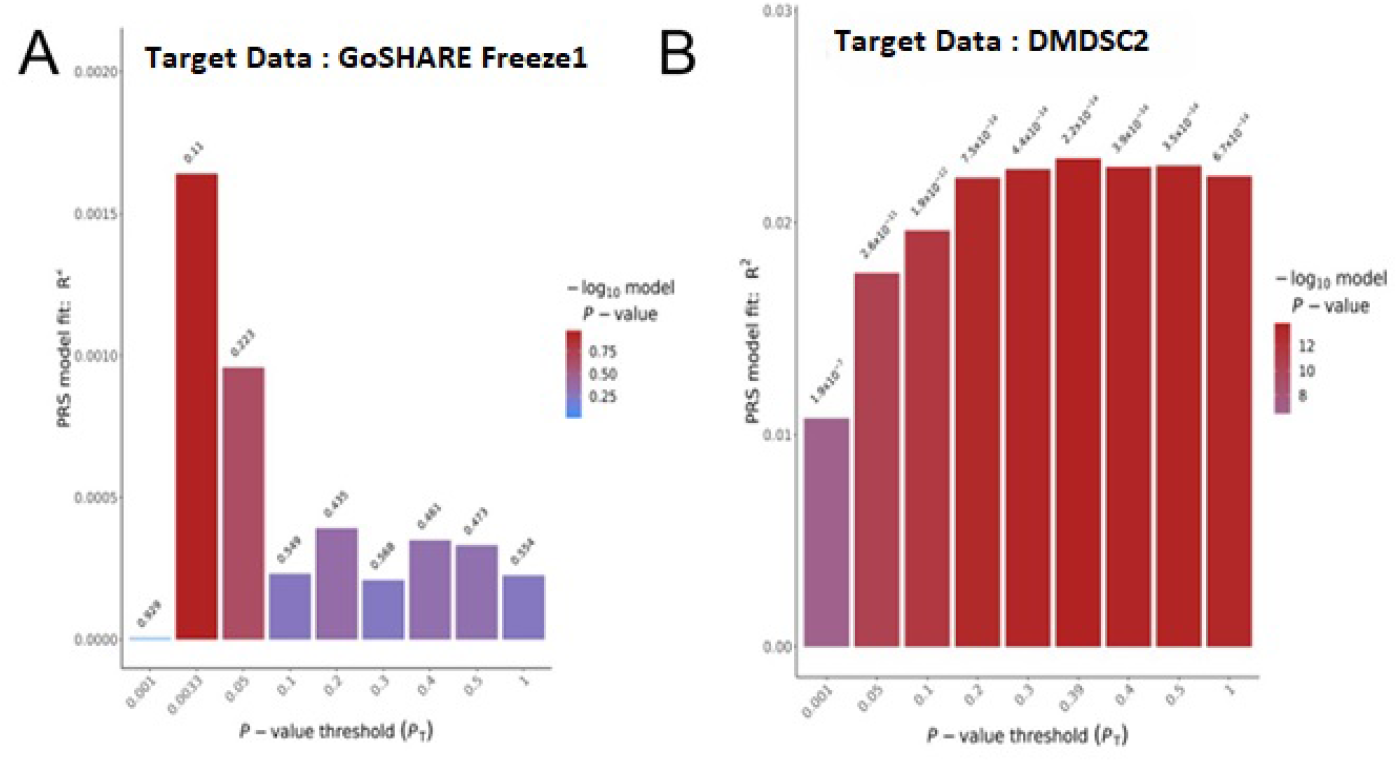
Performance of South Indian PRS in European population. PRS generated using Asian Indian summary data from DMDSC cohort 1 and tested in European samples (GoSHARE) and South Indian independent cohort (DMDSC 2) for polygenic risk prediction of age of onset.

## Discussion

In this study, we undertook a trans ethnic meta-analysis of age of diagnosis of T2D in 34,001 T2D individuals from two diverse ancestral backgrounds, European and Asian Indian, revealing a differential role for established T2D susceptibility loci in determining age of onset of diabetes. Interestingly the well-established T2D signal at *TCF7L2* was much more strongly associated with age of onset in the Asian Indian population compared to the European cohort, despite the allele frequency not differing between these ancestral groups. We showed that this difference was due to the distribution of age of onset of diabetes within the two ancestral groups, with the *TCF7L2* effect being largely observed in those diagnosed before the age of 50 in both ancestral groups. This is consistent with the concept that early onset disease would have a stronger genetic component; indeed, when we looked at the overall heritability estimates for age of diagnosis of T2D, the heritability was much stronger in the younger Asian Indian population with diabetes when compared to the more elderly European population with diabetes. We also found evidence for ethnic-specific signals that were associated with an early age at diagnosis of T2D in Asian Indians that were very rare in the European cohort. Our findings emphasize and support our recently reported finding that Asian Indians have greater genetic beta-cell dysfunction compared to Europeans[26].

The role of beta cell function as a driver for the early age onset of T2D in Asian Indians is well supported from our ethnic specific TCF7L2 and CDKAL1 signals. In addition, WDR 11 has previously been associated with fasting glucose[27], but not type 2 diabetes susceptibility per se.

One of the strengths of this study is that we address the lack of transferability and consistency of underlying age onset of T2D genetics between two ethnic groups. To date, this is the first study that demonstrates the genome wide PRS of age at onset T2D in Asian Indians. Overall, our polygenic risk scores findings derived from Asian Indian based GWAS can be useful for the population specific studies and it cannot be used to predict in Europeans where the genetics of T2D age onset itself is very distinct and unique between two ethnic backgrounds. We also noted that transferability of PRS across different ethnic groups demands careful evaluation.

One of the limitations in our study is the modest sample size of Asian Indian samples for the GWAS study, which limits our ability to identify associations with low-frequency variants. Next, our study cohort was limited only to the Asian Indian population and South Asian living in UK; thus, findings from the present study merits further validation in an independent larger discovery cohort. The biological interpretation of the significant Asian Indian T2D polygenic effects reported here needs further validation using an independent South Asian Indian cohort.

In conclusion, our study demonstrated the association of several previously established loci in European T2D GWAS for age onset of T2DM. However, we observed substantial heterogeneity in both the effect sizes and/or the allele frequencies between the ethnic groups. Furthermore, the higher heritability estimates of age of onset of type 2 diabetes in Asian Indians demonstrates the importance of further study of the genetic architecture of age of onset of type 2 diabetes in this ancestral group.

## Materials and Methods

### Ethics statement

All research has been conducted under the principles of the Declaration of Helsinki and approved by corresponding institutional review boards. All study participants provided written informed consent, and Research Ethics Committees approved the study.

### Study participants

We included participants from four independent cohorts: Dr Mohan’s Diabetes Specialties Centre (DMDSC), Chennai, India, Genetics of Diabetes Audit and Research in Tayside Scotland (GoDARTS), Genetics of Scottish Health Research Register (GoSHARE) and the United Kingdom Biobank (UKBB). DMDSC is a diabetes centre of single speciality hospitals and clinics established in 1991 in Chennai, Southern India, which includes currently there are 50 clinics in various locations across 10 states in India[28]. A total of 500,000 patients with type 2 diabetes to date, are provided with a unique identification number at their first visit, and clinical, anthropometric, and biochemical data are updated at each subsequent visit. Each patient underwent a comprehensive evaluation for screening and assessment of diabetes and presence of chronic complications at the time of their registered first visit, and these tests were repeated subsequently. All these data are collected and stored in the common diabetes electronic medical records (DEMR) system. GoDARTS consists of 18,306 participants from the Tayside region of Scotland, of which 10,149 participants were recruited based on their diagnosis of type 2 diabetes[16]. GoSHARE currently comprised of a biobank of around 74,000 individuals across NHS Fife, and NHS Tayside[17]. Both cohort participants’ have provided a sample of blood for genetic analysis and informed consent to link their genetic information to the anonymized electronic health records. UKBB is a large prospective general population cohort. A total of 502,628 individuals who were recruited 2006-2010 at age between 40 and 69 years from across the UK and provided electronically signed consent to use their self-reported answers on socio-demographic, lifestyle, ethnicity, a range of physical measures and blood or urine or saliva samples.

### Phenotyping – Age at Diagnosis of Type 2 Diabetes

We included 8,295 type 2 diabetes patients from DMDSC cohort of Asian Indians, whose first clinical visit was within one year of diagnosis. Diabetes is diagnosed by general practitioners using WHO criteria[29] for diagnosis and by the oral glucose tolerance test or fasting and/or random glucose test of HbA1c test. All study participants underwent a structured assessment, including detailed family history at the DMDSC. We excluded patients with type 1 diabetes or GADA positive. We selected the study participants in the GoDARTS, and GoSHARE based on the following inclusion criteria, aged between 20 and 80 years, the type 2 diabetes status diagnosed by general practitioners, and the clinical measurements as per WHO guidelines and recorded in the Scottish Care Information-Diabetes system. We identified 14,552 T2D participants within the UKBB cohort using diabetes diagnosis by the doctor (data-field code 2443), started insulin after a year diagnosis (2986) and the self-reported ethnicity (21000) and excluded participants with outlying principal components.

### Genotyping, Quality Controls, and Imputations

DMDSC genotyping was conducted for approximately 5,801 patients with type 2 diabetes by Illumina using the global screening arrays version (GSA v1.0); for the remaining participants (N=2,494) were genotyped on GSA v2.0. All genotyped samples were converted to PLINK format files using Illumina Genome Studio v2.04. We excluded samples if their call rate was less than 95% and genetically inferred sex discordance with phenotype data. We excluded the SNPs with less than 97 % call rate and HWE p-value less than 1e^-6^ (autosomal variants only). QC assessment was performed independently for DMDSC cohorts before and after phasing and imputation against the haplotype reference consortium (HRC r1.1) panel[30].

Genotyping of GoDARTS and SHARE cohorts derived from various platforms: Affymetrix 6.0 (Affymetrix, Santa Clara), Illumina Omni Express-12VI platform and GSA v.2.0, respectively. A total of 11,154 (6,999 GoDARTS and 4,155 GoSHARE) participants were considered after excluding those individuals failing to meet QC criteria. Individual genotype call rate (< 95%), heterozygosity>3 SD from the mean and the highly related sample’s identity by descent. We then carried out SNP-level QC by excluding markers < 97 % call rate, Hardy-Weinberg p< 1 × 10^-6^. Following sample and SNP quality control, Haplotype Reference Consortium (HRC) panel for both populations using the Michigan Imputation Server[31]. We retained all imputed SNPs with imputation information score > 0.4 and removed monomorphic markers.

The genome-wide genotyping for 488,377 participants in the UKBB was performed using custom-designed genotyping arrays including UK Biobank Axiom and UK BiLEVE Axiom Affymetrix array. We selected European and South Asian individuals in the UKBB based on PCA and self-reported ethnic background. A total of 14,552 participants fulfilled the phenotype criteria in this study. UK biobank genotyping, QC, PCA and imputation protocol are described elsewhere[32]. For the current study, we restricted the analyses to the UK Biobank participants who were in the full release imputed genomics datasets. We selected individuals of Asian Indians and European descents based on the principal component analysis (PCA) and the self-reported ancestry information.

### Ethnic-Specific Meta-analysis of GWAS

Genome-wide association analyses were performed independently for each cohort using an additive model while adjusting for sex (Fig 1). We estimated allelic effects using the using a linear mixed model as implemented in BOLT-LMM version 2.3.2[33] which accounts for relatedness and any population stratification and SNPTESTv2.5 in each cohort accordingly. We performed a meta-analysis based on ancestry: Asian Indians specific analyses include the DMDSC cohort, a unique Asian Indians representative data, and South Asians in the UKBB and the European specific analyses include the GoDARTS, GoSHARE, and White Europeans in the UKBB. There was no evidence of population stratification in meta-analysis (genomic inflation factor, λ = 1.007). We performed the meta-analyses using a fixed-effect method in METAL software, which assumes the effect allele is the same for each study within an ancestry. We then combined the summary statistics of GWAS from all the study populations. SNPs with imputation quality score < 0.40 were excluded from the analysis. Heterogeneity across these studies was assessed by the I^2^ (low to high) and Cochran’s Q statistics as reported by METAL. The Forest plot was generated using the metafor package[34]. We annotated the genetic variants using the University of California Santa Cruz (UCSC) Genome resource18 based on the Genome Reference Consortium Human genome build 37.

### Conditional analysis

We performed conditional analyses to identify additional secondary signals across the lead SNPs within the South Indian population.

### SNP-based heritability

We used the summary statistics data from the Asian Indians and Europeans specific meta-analyses to estimate the SNP-based heritability in a liability scale using Linkage Disequilibrium Score Regression (LDSC) software[35].

### Genome-wide Polygenic risk scores for age at diagnosis of type 2 diabetes

For the Polygenic risk scores (PRS), we considered summary statistics of GoSHARE and DMDSC samples. The PRSice tool generates the scores by the weighted sum of the risk allele carried by individuals based on effect estimate. We removed DNA polymorphisms with ambiguous strands (A/T or C/G) from the score derivation. SNPs were clumped to a more significant SNP in LD (r2 ≥ 0.10) within a 500 kb window. The PRS calculation considered several p-value thresholds (0.001, 0.05, and 0.1).

## Supporting Information

## Acknowledgements

This work was supported by the National Institute for Health Research using Official Development Assistance (ODA) funding [INSPIRED 16/136/102]. The Wellcome Trust United Kingdom Type 2 Diabetes Case-Control Collection (supporting GoDARTS) was funded by The Wellcome Trust (072960/Z/03/Z, 084726/Z/08/Z, 084727/Z/08/Z, 085475/Z/08/Z, 085475/B/08/Z) and as part of the EU IMI-SUMMIT program. The present study was conducted using the UK Biobank Resource under application No. 20405. We are thankful to all the families who took part in this study. We are grateful to the GoDARTS, SHARE and DMDSC teams, including interviewers, computer and laboratory technicians, clerical workers, research scientists, volunteers, managers, receptionists, healthcare assistants, and nurses, for their cooperation in recruiting them. We would like to acknowledge Dundee Health Informatics Centre (HIC) for managing and providing anonymised data.

## Author Contributions

Study concept and design: CNAP.

Data collection and access: GoDARTS, GoSHARE, UKBB - CNAP, AD, ERP

Data collection and access: DMDSC - MV, ARM, RV

Data processing: MKS, SS, SL, NS.

Data analysis: SS.

Drafting of the manuscript: SS and CNAP.

Critical revision of the manuscript: All authors involved in the critical revision and approved the final version for publication.

